# The Impact of Host Genetics and/or its Microbiota on the Severity of *Corynebacterium*-Associated Hyperkeratosis in Outbred Athymic Nude Mice

**DOI:** 10.1101/2024.10.06.616361

**Authors:** Abigail Michelson, Christopher Cheleuitte-Nieves, Ileana C. Miranda, Irina Dobtsis, Neil S Lipman

## Abstract

*Corynebacterium bovis* (Cb), the etiology of *Corynebacterium*-associated hyperkeratosis (CAH) in nude mice, may impact research outcomes. Little is known about the differences in the course and severity of CAH in different outbred athymic nude mice stocks. Three genetic stocks (designated A, B, and C), 1 of which was obtained from 2 geographically separate colonies with distinct microbiota (A1 and A2), were inoculated topically with 1 × 10^8^ CFUs of a pathogenic Cb field isolate (#7894; n = 6/stock) or sterile media (n=2/stock; controls). Clinical signs were assessed daily and scored 0 – 5 based on lesion severity. Mice were euthanized at 14 (A1, A2, B, and C) or 28 (B) days-post-inoculation (dpi), macroscopic changes documented, and 6 skin samples per mouse were obtained and histologically scored 0 – 4 based on the presence and severity of hyperkeratosis, acanthosis, inflammation, and bacterial colonies. No stock A1 or control mice developed clinical disease; 1 of 6 stock B mice developed mild CAH (mean peak clinical score [MPCS] – 0.33) at 14 dpi (14 day group) and 2 of 6 stock B (28 day group) developed mild CAH at 15 dpi (MPCS -0.33); and, all stock C and A2 mice developed significant clinical signs at 5 dpi (MPCS – 2.5 and 3, respectively) which resolved by 11 dpi. Despite differences in clinical presentation, all Cb-infected mice had hyperkeratosis and/or acanthosis with associated bacterial colonies. Stocks A1 and B, which had minimal or no clinical signs, were colonized with *Corynebacterium amycolatum* (Ca). In contrast, stocks C and A2 were not colonized with Ca, raising the possibility that Ca and/or other components of the skin microbiota may mitigate clinical signs but not necessarily all histopathologic changes associated with infection. These findings suggest host genetics and/or the skin microbiota can markedly influence the presentation of CAH in nude mice.

## Introduction

Corynebacterium-associated hyperkeratosis (CAH), also known as scaly skin disease, results following infection with the Gram-positive bacterial rod, *Corynebacterium bovis* (Cb). The bacterium colonizes the skin of immunocompromised mice, significantly impacting research in which infected mice are used.^1, 12, 18^ Cb infection results in 100% morbidity with, in general, low mortality in susceptible, immunocompromised strains in which it causes a diffuse scaling dermatitis and alopecia (in hirsute strains), and acanthosis and orthokeratotic hyperkeratosis microscopically.^1,6,19^ Nude mice typically present with clinical signs 7-10 days post infection which can include in addition to the dermatopathology, weight loss, dehydration, and pruritis.^6,7,12,16^ Variable success in the management and eradication of Cb has been achieved through antibiotic therapy, colony depopulation, and decontamination.^1,10,12,13^ However, once infected, immunocompromised mice remain colonized with Cb despite antibiotic treatment and in some cases, clinical resolution.^1,10,12^ Antibiotic use can also confound research by altering the anti-tumor response to chemo- and immunotherapy as well as the microbiota, shifting the host immune response against pathogens, as examples.^9,17^

Surprisingly, an outbred nude mouse stock used in a pilot study investigating a novel therapeutic strategy for Cb failed to develop clinical disease when challenged with a Cb isolate and dose that was previously shown to be highly pathogenic in another nude mouse stock.^12^ In addition to the recognized genetic differences in the 2 stocks, the mice used in the pilot study also included *Corynebacterium amycolatum* (Ca) as a component of their skin microbiota.

A study was undertaken to ascertain whether the resistance to Cb-associated clinical disease was a result of the skin microbiota of the nude mouse stock utilized. Three outbred athymic female nude mouse stocks, 2 of which were colonized with Ca, 1 of which was available from 2 distinct colonies with different skin microbiota (1 of which included the presence of Ca), were challenged with a high dose of a pathogenic Cb isolate (#7894) and then evaluated for colonization; clinical disease development, progression, and resolution; and histopathologic changes.

## Materials and Methods

### Experimental design

Three genetic stocks of athymic female nude mice (n=6 stock or colony), each from a distinct vendor (designated A, B and C), 1 of which was obtained from 2 geographically separate colonies with presumed distinct microbiota (A1 and A2), were inoculated topically with 1 × 10^8^ CFUs of a pathogenic Cb field isolate (#7894).^5,12^ Two additional mice of each stock and/or colony were inoculated similarly with sterile bacterial media to serve as negative controls. Clinical signs were assessed daily and scored 0 – 5 based on lesion severity (See Table 1). Cb colonization was assessed weekly (1 week pre-inoculation, on the day after inoculation, and every 7 days-post-inoculation [dpi]). Mice were euthanized by CO_2_ overdose at either 14 (A1, A2, B, and C) or 28 (B) dpi, a gross necropsy was performed, macroscopic changes documented, and 6 full-thickness skin samples per mouse were obtained, assessed histologically, and scored. Stock B mice were assessed at 2 post-inoculation timepoints (14 and 28 days; n = 6/timepoint). This decision was based on a prior assessment in which the stock’s skin microbiota, which includes *Corynebacterium amycolatum* (Ca), appeared to protect the mice from the development of Cb-associated skin disease. The extended assessment aimed to determine if Ca delayed development of Cb-associated disease. Similarly, stock A mice were obtained from 2 distinct colonies as the colonies differed in that mice from A1 skin microbiota includes Ca.

**Table 1:** *C. bovis* clinical scoring system for nude mice. Animals were graded 0-5 based on clinical signs representing mild, moderate, and severe disease.

| Grade | Clinical signs |
| --- | --- |
| 0 | No clinical signs observed. |
| 1 | Mild amount of flakes/scales adhered to the skin and/or mild hunched posture with no erythema. |
| 2 | Mild amount of flakes/scales adhered to skin and/or mild hunched posture and mild erythema associated with flaking/scaly lesions. |
| 3 | Moderate erythema and/or moderate amount of white flakes adherent to the skin, or minimal white flakes on 3 or more additional locations affected including the ventrum, muzzle, cheeks, crown, and limbs. |
| 4 | Moderate erythema and/or moderate amount of white flakes adherent to the skin and moderate hunched posture. |
| 5 | Severe amount of flakes/scales adhered to skin (thick layer), and/or severe erythema, and/or severe hunched posture. |

### Animals

Female, 5 to 9-week-old, outbred athymic nude mice (Crl:NU(NCr)-*Foxn1*^*nu*^, Charles River Laboratories, Wilmington, MA; Hsd:Athymic Nude-*Fox1*^*nu*^, Inotiv, Livermore, CA and Indianapolis, IN; Jax:NU/J, The Jackson Laboratory, Bar Harbor, ME; n=8/stock and/or site). One genetic stock (Hsd:Athymic Nude-*Fox1*^*nu*^) was obtained from 2 distinct geographic colonies based on their microbiota as mice from 1 of the 2 sites are colonized with Ca. Mice were confirmed free of Cb and Ca (unless otherwise noted) by aerobic culture on arrival. Mice were free of the following agents: Sendai virus, Pneumonia virus of mice, mouse hepatitis virus, minute virus of mice, mouse parvovirus, Theiler meningoencephalitis virus, reovirus type 3, epizootic diarrhea of infant mice (mouse rotavirus), murine adenovirus, murine astrovirus 2, polyoma virus, K virus, murine cytomegalovirus, mouse thymic virus, lymphocytic choriomeningitis virus, Hantaan virus, ectromelia virus, lactate dehydrogenase elevating virus, mouse norovirus, and murine Chaphamaparvovirus, *Bordetella bronchiseptica, Citrobacter rodentium, Clostridium piliforme, Corynebacterium bovis, Corynebacterium kutscheri, Filobacterium rodentium, Mycoplasma pulmonis, Chlamydia muridarum, Salmonella spp*., *Streptobacillus moniliformis, Encephalitozoon cuniculi*, ectoparasites, endoparasites, and enteric protozoa.

### Husbandry and housing

Mice were housed in autoclaved, solid bottom and top, gasketed and sealed, polysulfone, positive-pressure, individually ventilated cages (Isocage, Allentown Caging Equipment Company, Allentown, NJ) on autoclaved aspen chip bedding (PWI Industries, Quebec, Ontario, Canada) in distinct cages based on whether the mice were experimental or control, as well as by stock and/or site. Each cage was ventilated with HEPA-filtered air (filtration at rack and cage level) at approximately 30 air changes per hour and the HEPA-filtered rack effluent was exhausted directly into the building’s HVAC system. Mice received gamma-irradiated, autoclaved feed (5KA1, Purina Mills International, St Louis, MO) and autoclaved acidified (HCl) reverse-osmosis–purified water (pH 2.5 to 2.8) ad libitum. Each cage was provided with a sterile bag of Glatfelter paper containing 6g of crinkled paper strips (EnviroPAK, WF Fisher and Son, Branchburg, NJ) for enrichment. Cages were changed every 2 weeks, and all manipulations were performed in a class II, type A2 biologic safety cabinet (BSC; LabGard S602-500, Nuaire, Plymouth, MN) using aseptic technique with sterile gloves. Control mice were changed before experimental animals to reduce the likelihood of cross contamination. The animal holding room was ventilated with 100% fresh air at a minimum of 10 air changes hourly and maintained at 72±2°F (21.5±1°C), relative humidity between 30% and 70%, and a 12:12-h light: dark photoperiod (lights on at 0600, off at 1800). All animal use was approved by Memorial Sloan Kettering’s (MSK’s) IACUC and conducted in accordance with AALAS’s position statements on the Humane Care and Use of Laboratory Animals and Alleviating Pain and Distress in Laboratory Animals. MSK’s animal care and use program is AAALAC-accredited and operates in accordance with the recommendations provided in the Guide for the Care and Use of Laboratory Animals (8^th^ edition).^8^

### *Corynebacterium bovis* propagation and inoculation

Cb isolate #7894 stored as a frozen stock was grown on trypticase soy agar supplemented with 5% sheep blood (BBL TSA II 5% SB, Becton Dickinson, Sparks, MD) at 37°C with 5% CO_2_ for 48h, as previously described.^3,5^ The growth curve for this isolate had been established.^5^ The bacterial inoculum was collected from the culture during mid-log of the exponential growth phase, titrated and serially diluted in brain heart infusion broth (BHI; Becton Dickinson) supplemented with 0.1% Tween 80 (VWR Chemicals, Solon, OH) to obtain 10^8^ +/-15% bacteria/50 µl as previously described.^12^ The Cb inoculum was placed in sterile 2 ml polypropylene centrifuge tubes (Fisher Scientific, Waltham, MA), transferred to the vivarium on ice and the animals were inoculated within an hour of inoculum preparation. Inoculation was performed in a class II, type A2 BSC (LabGard S602-500, Nuaire) using sterile materials and aseptic methods by personnel donning sterile gloves. After inoculation, the concentration of each inoculum was confirmed and enumerated by plate count.

For inoculation and subsequent sample collection for aerobic bacterial culture, control mice were handled before experimental animals. Each cage containing an experimentally naïve animal was removed from the rack and all sides of the cage were liberally sprayed with disinfectant (Peroxigard [1:16], Virox Technologies Inc., Oakville, ON) and placed in the BSC. Each cage was opened for less than 2 minutes. Bacterial inoculation was performed with the mouse restrained with the handler’s non-dominant hand by gently grasping the tail base and the animal’s hindquarters were elevated slightly while the animal grasped the wire bar lid with its forelimbs. The lid was positioned to allow the restrained animal to remain in the horizontal plane. Using the dominant hand, the bacterial inoculum was applied directly to the skin using a sterile filter micropipette (P200N, Marshal Scientific, Hampton, NH) along the dorsal midline starting at the nuchal crest proceeding to the tail, a distance of ∼ 2 cm. A 50 μl bacterial suspension was evenly distributed along the dorsal midline at 4-5 sites dispensing 10-15 μL per site using a sterile filtered 200 uL pipette tip (Filtered Pipet Tips, Crystalgen, Commack, NY). This method ensured that the bacterial suspension did not flow off the animal. The cage was closed and sprayed thoroughly with disinfectant prior to returning it to the rack. All interior surfaces of the BSC were sprayed with disinfectant and a new pair of sterile gloves were donned between each cage. A new cage was placed in the BSC as previously described and the previous steps repeated until all inoculations (0 or 10^8^ bacteria/mouse) were completed. Animals in the control group were inoculated with an equivalent amount of bacterial-free BHI broth.

### Bacterial sampling and culture

All mice were confirmed negative for Cb and Ca by aerobic culture prior to commencing the study except sites A1 and B which were negative for Cb and positive Ca. Cb infection was assessed weekly by aerobic skin culture for up to 28 dpi (B only). To prevent cross-contamination within each dose group, controls followed by animals with lower clinical scores were swabbed first. When animals had similar scores, cages were selected at random. All cages were handled aseptically as previously described. Each cage was opened for no longer than 5 min. The animal was grasped gently by the base of the tail, allowing all 4 limbs to grasp the wire bar lid. A sterile culture swab (BBL CultureSwab, Becton Dickinson) was held perpendicular to the animal and swabbed along the dorsal midline from the base of the tail cranially to the nuchal crest and caudally back to the tail base, with rotating of the swab as it was advanced. The cage was closed, sprayed, and wiped down with disinfectant and returned to the rack. The BSC was cleaned as described above, the handler donned a new pair of sterile gloves, and the next cage was processed in the same manner as above.

Skin swabs were each streaked in a 4-quadrant pattern onto agar plates (TSA II 5% SB, Becton Dickinson).^2^ The plates were incubated for 72h at 37°C with 5% CO_2_. When present, distinct colonies were speciated by using MALDI-TOF spectroscopy (MALDI Biotyper Sirius CA System, Bruker, Billerica, MA). Plates without growth were held for an additional 7 days before being considered negative. Plates were scored from + to ++++ based on the number of quadrants with bacterial growth.

### Clinical scoring

Mice were assessed daily for either 14 or 28 dpi. Animals were evaluated cage side and scored using the system utilized by Mendoza et al. and provided in Table 1.^12^

### Postmortem gross examination and skin histopathology

On 14 or 28 dpi, mice were euthanized by CO_2_ overdose. Each mouse underwent a complete postmortem gross examination, including macroscopic inspection of the skin and all internal organs. Full-thickness skin samples (n=6), approximately 1 cm in length, were collected from the ear, head (between the ears), dorsum and ventrum (centromedial), and fore- and hindlimb (anterior) using a scalpel from each mouse. Specimens were fixed in 10% neutral buffered formalin, processed in alcohol and xylene, embedded in paraffin, sectioned at 5µm, and stained with hematoxylin and eosin. Histologic examination was performed on all skin biopsies to confirm the presence or absence of Cb-related lesions and to determine the magnitude of the histologic changes. Slides were evaluated and lesions recorded by a board-certified veterinary pathologist (ICM) who was blind to groups. Samples were semi-quantitatively scored as normal (0), minimal (1), mild (2), moderate (3), or severe (4), depending on the degree of acanthosis, orthokeratotic hyperkeratosis (orthokeratosis), bacteria, and inflammation.^12^ Specifically, acanthosis refers to an elevated number of keratinocytes and indicates an increase in epidermal thickness resulting from hyperplasia and often hypertrophy of cells of the stratum spinosum. Minimal acanthosis was defined as an increased thickness of up to a 3-cell stratum spinosum layer; mild as 4 to 7 cell layers; moderate as 8 to 10 cell layers; and, severe as greater than 10 cell layers. Orthokeratosis refers to increased thickness of the stratum corneum without retention of keratinocyte nuclei and was mostly identified in a compact to laminated pattern. Minimal orthokeratosis was defined as an increased thickness of up to 50µm; mild as 100 to 200µm; moderate as to 200 to 300µm; and, severe as greater than 300µm. Bacterial colonies were assessed based on the presence of distinguishable clusters of short bacilli in the stratum corneum or the lumen of hair follicles. Minimal presence of bacteria was defined as small amounts of scattered bacterial rods with no formation of characteristic clusters; mild as 1 to 2 clusters; moderate as 3 to 5 clusters; and, severe as greater than 6 clusters. Inflammation was mostly comprised of a mixture of mononuclear cells and neutrophils, associated both with and without hair follicle rupture. Minimal inflammation was defined as single superficial pustules or small numbers of inflammatory cells scattered in the dermis; mild as slightly larger or increased numbers of superficial pustules or slightly greater numbers of inflammatory cells scattered in the dermis; moderate as larger or increased numbers of pustules or increased numbers of inflammatory cells in the dermis, frequently forming clusters around capillaries or adnexal units; and, severe as large and multiple pustules with high numbers of inflammatory cells forming coalescing clusters or sheets in the dermis, often infiltrating the epidermis or adnexal epithelia. For all stocks and sites, the average of each of the 4 microscopic characteristics was calculated using 6 biopsies per mouse. The “total histopathology score” was calculated by adding the average of all 4 histologic characteristics. Additionally, to ascertain whether the Cb-induced pathologic changes differed across the 6 sites that were assessed, the site-specific histopathology scores were compared by averaging the scores from all the Cb-infected stocks.

### Statistical analysis

Mean peak clinical scores between Ca+ and Ca negative stocks and mean daily clinical and mean total histopathology scores across stocks and/or sites were compared. Difference in mean peak clinical scores were compared using unpaired t and Mann-Whitney U tests. Differences in daily clinical scores across stocks and/or sites were compared using one-way ANOVA, Kruskal-Wallis and Dunnet’s multiple comparison tests. Differences in duration of clinical presentation between stocks and sites were compared using one-way ANOVA, Kruskal-Wallis and Dunnet’s multiple comparison tests. Differences in histologic characteristics (including hyperkeratosis, acanthosis, bacterial colonies, and inflammation) were compared between stocks and sites, and their respective controls using unpaired t and Mann-Whitney U tests. The mean of histopathology scores of the 6 sections of skin for each histopathologic criteria assessed for each of the stocks and/or geographical site were also compared using a one-way ANOVA, Kruskal-Wallis and Dunnet’s multiple comparison tests. P values <0.05 were considered statistically significant. All calculations were performed using Prism 10 for Windows (GraphPad Software version 10.3.0, San Diego, CA).

## Results

### Clinical score and disease progression by stock and colony location

The temporal course of clinical disease, which differed by outbred athymic nude stock (stock A, B and C) and by the geographical site for stock A mice (A1 and A2), is shown in Figure 1. A2 and C nude mouse stocks had a very similar clinical course with skin lesions first detected on 5 dpi progressing and reaching mean peak clinical score (MPCS +/-SD) of 2.5 +/-0.5 (A2) or 3 +/-0.58 (C) on 7 and 8 dpi, respectively, with observable lesions resolving by 11 dpi. Mice from site A1 never developed observable skin lesions, while stock B mice did not develop observable lesions until 14 (14-day study) or 15 (28-day study) dpi with MPCSs of 0.33 +/-0.75 (14-day study) and 0.33 +/-0.47 (28-day study), respectively. Comparing differences at specific timepoints, the mean clinical scores for stocks A2 and C were significantly higher than the scores assigned to stocks A1 and B (for both the 14- and 28-day study groups on days 5, 6, 7, and 9 post infection, whereas on day 8 only stock C’s score was significantly higher.

While all stock A2 and C mice developed skin lesions, only 1 of 6 (14-day study) and 2 of 6 (28-day study) stock B mice developed skin lesions. The duration of detectable lesions ranged from as short as 0 days in stock A1 mice to as long as 6 days for stocks A2, B, and C, with durations (mean +/-SD) and number of mice per stock out of 6 presenting with clinical signs of 5.2 +/-0.4; 6/6 (A2), 4.5 +/-2.1; 2/6 (B 28 dpi), and 5.5 +/-0.55; 6/6 (C) days. The duration of clinical presentation for the mice that did demonstrate detectable lesions was compared between stocks, but there were no statistically significant differences (data not shown).

Interestingly, stocks of nude mice which included Ca as a component of their skin microbiota had a significantly lower mean clinical score (0.22 +/-0.53) than mice that were not colonized with Ca (2.75 +/-0.6; Figure 2). None of the control mice of any stock developed clinical signs.

**Figure 1:**
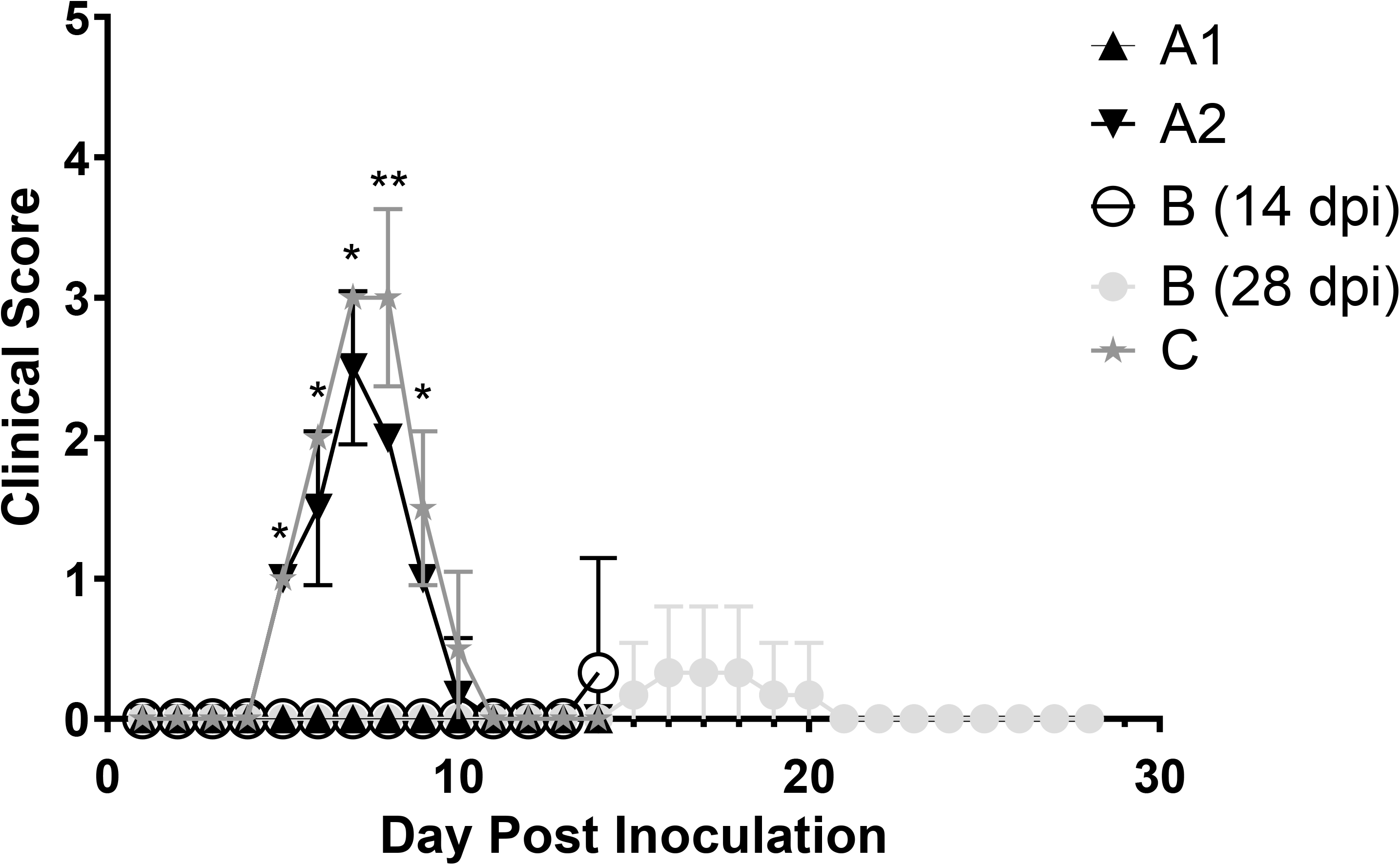
Clinical scores by stock and colony geographical site. Scores reflect the mean +/-SD for n=6 mice/stock assessed daily. Stock A mice were obtained from 2 geographical sites with mice from A1 colonized with Ca while A2 were Ca-free. Stock B mice were evaluated for either 14 or 28 dpi. *p<0.05 when comparing stock A2 and C to A1 and B (14- and 28-day study). *\*\** p<0.05 when comparing stock C to stocks A1 and B (14- and 28-day study).

**Figure 2:**
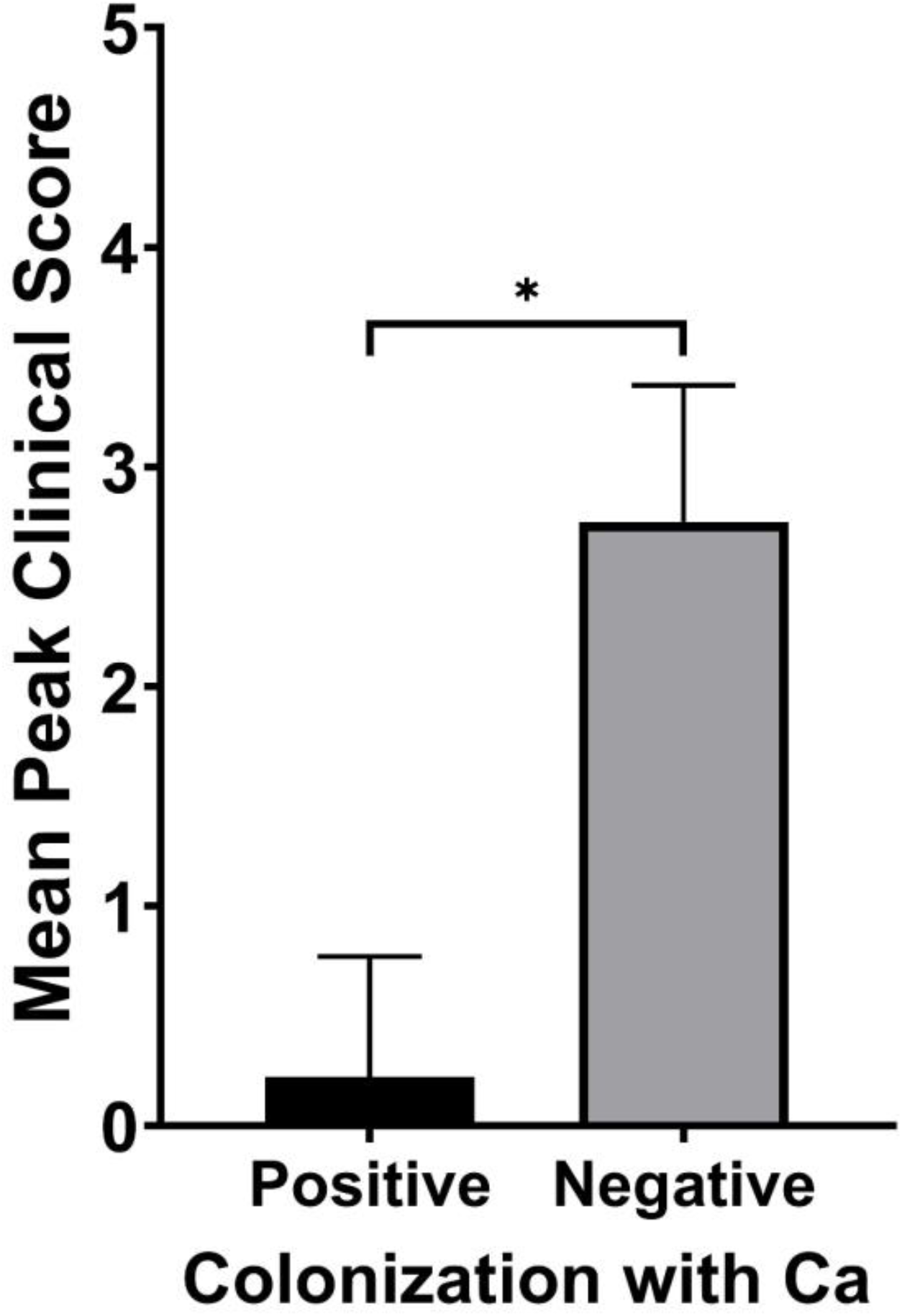
Stocks with Ca as part of the normal skin microbiota had a statistically significant higher mean peak clinical score than those without Ca (p<0.05).

### Bacterial culture

Semiquantitative bacterial colonization was assessed weekly throughout the course of the study (Tables 2A and B). Stocks A1 and B were Ca culture positive on intake, while stocks A2 and C were negative. The Ca positive mice (A1 and B) were all culture positive (+++) on intake, but ultimately became Ca culture negative. Stock A1 mice became Ca culture negative 7 (Cb-infected) or 14 (controls) dpi. In contrast, stock B mice became culture negative 21 dpi, except Ca (+) was isolated from the control group at 28 dpi. All mice inoculated with Cb were culture positive (+++) at most time points post inoculation except for B week 2 (+) and C week 1 (+).

**Table 2.** A. Semiquantitative assessment of skin colonization with *C. amycolatum*.

| Stock/Site | Intake | Week 1 | Week 2 | Week 3 | Week 4 |
| --- | --- | --- | --- | --- | --- |
| **A1 controls** | +++ | + | - | NA | NA |
| **A1 exp** | +++ | - | - | NA | NA |
| **B controls** | +++ | + | ++ | - | + |
| **B exp** | +++ | +++ | + | - | - |
| **A2 and C (exp and controls)** | - | - | - | NA | NA |

B. Semiquantitative assessment of skin colonization with *C. bovis*.
| Stock/Site | Intake | Week 1 | Week 2 | Week 3 | Week 4 |
| --- | --- | --- | --- | --- | --- |
| **A1** | - | +++ | +++ | NA | NA |
| **A2** | - | +++ | +++ | NA | NA |
| **B** | - | +++ | + | +++ | +++ |
| **C** | - | + | +++ | NA | NA |
| **Controls**<br>**A1, A2, B & C** | - | - | - | - (B only) | - (B only) |

NA, not applicable.

### Histopathology

Skin pathology was assessed 14 or 28 dpi at which point all mice, except for 1 mouse from stock B assessed at 14 dpi, had no clinically observable lesions. The cumulative histopathology scores of all Cb-infected mice were roughly 3-to 5-fold higher than their respective controls, although these differences were only significant for stock B (Figure 3 and Table 3). However, all individual components of the scoring system were significantly higher in the Cb-infected mice across stocks, except for the degree of bacterial colonies for stock B at 14 dpi. (Figure 3 and Table 3). All mice, including controls, had inflammation; however, the magnitude of inflammation, although relatively mild, was statistically more severe in Cb-infected mice. None of the controls had hyperkeratosis and/or observable bacteria, both of which were consistently observed in Cb-infected mice. While detected in all Cb-infected mice, acanthosis was inconsistently observed in the controls. The cumulative score in stock A1 mice was significantly different from stocks B and C (Figure 3 and Table 3). There were minimal differences in scores from stock B mice when comparing mice sacrificed at 14 (3.25 +/-0.52) versus 28 (3.28 +/-1.11) dpi.

**Figure 3:**
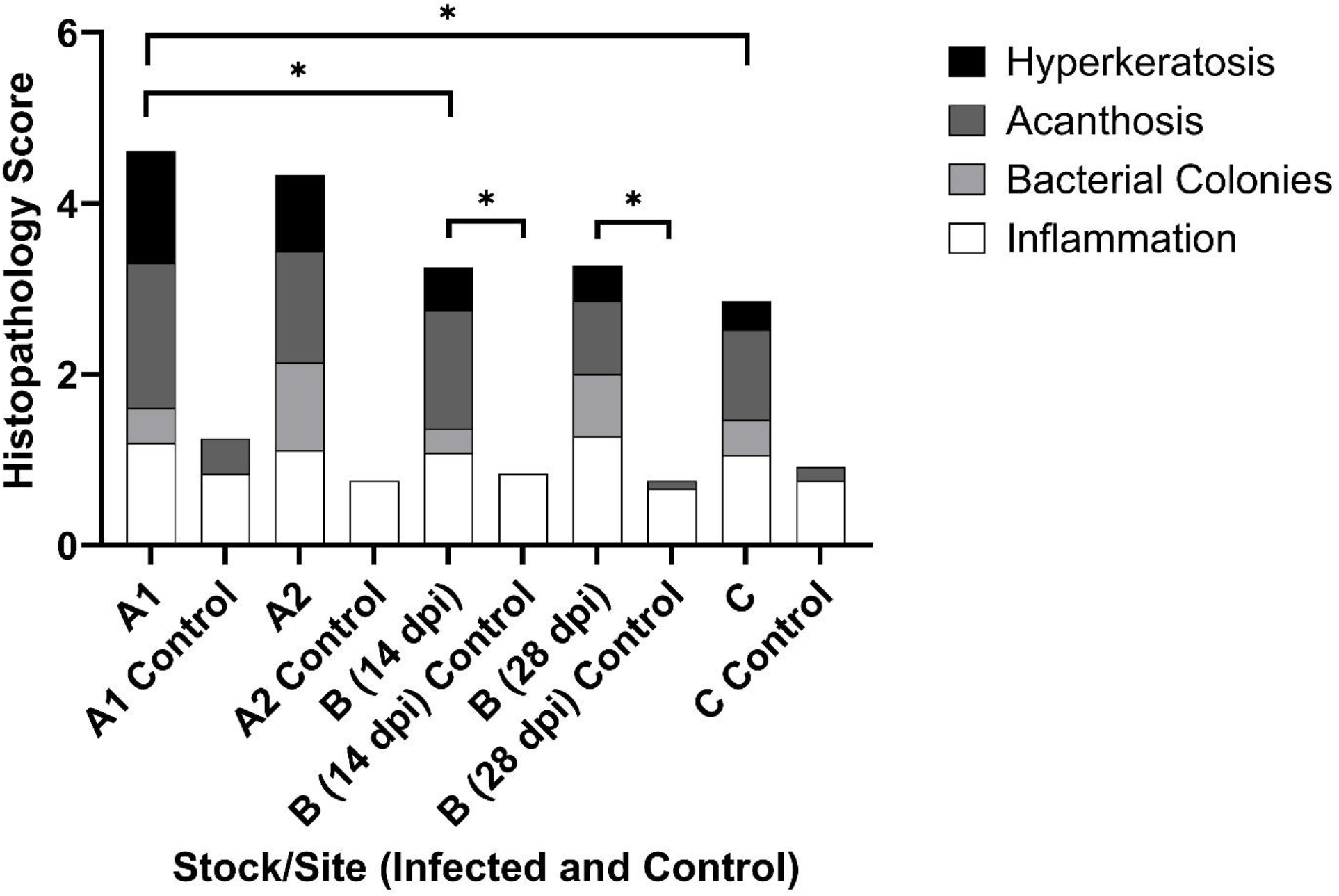
Graphical presentation of histopathology scores for 3 Cb-infected and uninfected control outbred nude stocks (A, B and C) and geographical sites (A1 and A2) evaluated 2 and/or 4 (B only) weeks post-inoculation. Scores reflect the average distribution of the means (6 biopsies/mouse) of the 6 Cb-infected or 2 uninfected control mice/stock for each of 4 histopathology criteria assessed. *p<0.05.

**Table 3:** Histopathology scores for all stocks and sites assessed.

| Stock/Site | A1 |  |  |  |  | A2 |  |  |  |  | B (14 dpi) |  |  |  |  | B (28 dpi) |  |  |  |  | C |  |  |  |  |
| --- | --- | --- | --- | --- | --- | --- | --- | --- | --- | --- | --- | --- | --- | --- | --- | --- | --- | --- | --- | --- | --- | --- | --- | --- | --- |
| Histopathology Criteria | H | A | B | I | T | H | A | B | I | T | H | A | B | I | T | H | A | B | I | T | H | A | B | I | T |
| Control | 0 | 0.42<br>+/-<br>0.67 | 0 | 0.83<br>+/-<br>0.39 | 1.25<br>+/-<br>0.12 | 0 | 0 | 0 | 0.75<br>+/-<br>0.45 | 0.75<br>+/-<br>0.12 | 0 | 0 | 0 | 0.83<br>+/-<br>0.39 | 0.83<br>+/-<br>0 | 0 | 0.08<br>+/-<br>0.29 | 0 | 0.67<br>+/-<br>0.49 | 0.75<br>+/-<br>0.35 | 0 | 0.17<br>+/-<br>0.39 | 0 | 0.75<br>+/-<br>0.45 | 0.92<br>+/-<br>0.12 |
| Experimental | **1.31**<br>+/-<br>**0.47** | **1.7**<br>+/-<br>**0.53** | **0.42**<br>+/-<br>**0.6** | **1.19**<br>+/-<br>**0.4** | 4.61<br>+/-<br>0.57 | **0.89**<br>+/-<br>**0.67** | **1.31**<br>+/-<br>**0.66** | **1.03**<br>+/-<br>**0.97** | **1.11**<br>+/-<br>**0.32** | 4.33<br>+/-<br>0.37 | **0.5**<br>+/-<br>**0.56** | **1.39**<br>+/-<br>**0.73** | 0.28<br>+/-<br>0.51 | **1.08**<br>+/-<br>**0.28** | **3.25**<br>+/-<br>**0.52** | **0.42**<br>+/-<br>**0.55** | **0.86**<br>+/-<br>**0.87** | **0.72**<br>+/-<br>**0.74** | **1.28**<br>+/-<br>**0.45** | **3.28**<br>+/-<br>**1.11** | **0.33**<br>+/-<br>**0.48** | **1.06**<br>+/-<br>**0.58** | **0.42**<br>+/-<br>**0.55** | **1.06**<br>+/-<br>**0.23** | 2.86<br>+/-<br>0.88 |

Scores reflect the mean +/- SD for each assessed histologic feature. Bolded scores are statistically significant as compared to their respective controls ( $p \leq 0.05$ ). H, hyperkeratosis; A, acanthosis; B; bacterial colonies; I, Inflammation; T, cumulative histology score (H + A + B + I = T).

To ascertain whether the Cb-induced pathologic changes differed across the 6 anatomical sites that were assessed, the histopathology scores were compared. There were no significant differences appreciated across sites (data not shown).

## Discussion

This study demonstrates for the first time that the magnitude of clinical disease following infection with Cb can be influenced by the genetic stock and colony source of nude mice. Two of the stocks of outbred nude mice assessed failed to develop or developed minimal clinical disease following experimental infection with a Cb isolate and dose that was previously shown, using 1 of the stocks evaluated in this study (A2), to be the most pathogenic of the isolates evaluated in the prior study.^12^ This finding may have practical implications for enzootically infected colonies in which control of Cb is logistically challenging.^3^ It is plausible that the replacement of a ‘susceptible’ with a ‘resistant’ nude mouse stock could reduce or eliminate the development of clinical signs following Cb infection and over time, the magnitude of the Cb environmental burden may be reduced as potentially there would be fewer mice shedding the bacterium and those that shed would be shedding fewer bacteria. While this result is plausible, it is important to recognize that only a single Cb isolate was evaluated in this study and the differences in sensitivity to this isolate may not translate to other isolate(s) circulating in a colony. Additionally, changing stock may be scientifically problematic as the genetic and microbiota differences between stocks could confound research findings as there may be differences in the animals’ response to the experimental variable being studied or the growth characteristics of the engrafted tumor, for which nude mice are typically employed, may be altered.

Interestingly, the presence of Ca, either alone and/or in combination with other microorganisms, appears to mitigate the clinical presentation of CAH as most mice utilized in this study that were colonized with Ca did not present clinical signs following experimental inoculation with Cb #7894. Additionally, of the Ca+ mice that developed clinical disease, their mean peak clinical scores were significantly lower than observed in Ca-free mice. The duration of visible lesions was also shorter in Ca+ mice, but this difference was not statistically significant. This lack of significance likely reflects the limitation of the study’s statistical power. A larger group size could have yielded significance. To ascertain whether Ca simply delayed Cb colonization and disease, which has been shown to occur in NSG mice, a group of stock B mice were monitored for a longer 28-day course which did reveal a delayed CAH presentation in 2 mice, although their clinical scores were quite low.^10,11,12^ These findings suggest that colonization with Ca and/or other unidentified microorganism(s) may provide some degree of protection against development of CAH and its’ severity. The mechanism of protection may be due to competitive exclusion, a condition by which two bacterial species occupying the same environment and relying on similar resources cannot equally coexist.^4^ This theory could explain either or both the decreased severity as well as the delay in onset of CAH in Ca+ mice. Further investigation is warranted to better understand the role of Ca and the microbiota on Cb colonization and Cb-induced disease.

It would be intriguing to determine whether host genetics also plays a role in the resistance to Cb-induced disease and colonization as all 3 of the outbred nude mouse stocks used in this study are distinct outbred stocks which would be expected to have genetic differences. Are stock B mice more resistant to Cb as a result of colonization with Ca and/or other microbiota components and/or their genotype? A study in which each of the stock’s microbiota would be established in each of the 3 stocks and the resulting 9 possible stock/microbiota combinations subsequently challenged with the same Cb isolate would provide further clarity as to the role of Ca, the microbiota, and/or the host’s genetics has on the development of CAH.

Unfortunately, the semi-quantitative aerobic culturing method used in this investigation cannot adequately reveal the dermal population dynamics of Ca and Cb. Ca colonization appears to diminish temporally, irrespective of infection with Cb, perhaps negating the competitive exclusion theory that Ca outcompetes Cb. For mice that were Ca+ and Cb-infected, the temporal decrease in Ca colonization could be a result of Cb outcompeting Ca, but this does not explain why there is a reduction of Ca in the uninfected controls nor why CAH would be mitigated. Conversely, once the Ca+ mice were infected with Cb, colonization with Cb persisted but the degree of colonization varied slightly among stocks. Ca may mitigate or may simply serve as a marker for other dermal microbial inhabitants which are the primary influencers of Cb disease resistance. In humans, studies evaluating the skin microbiota have documented numerous challenges, including its irregular distribution, the instability of some dermal microorganisms, and the impact of the external environment.^14^ There are some studies that have evaluated the skin microbiota of laboratory and wild mice, but none have evaluated temporal variability, which is a known characteristic of the microbiota of human skin.^1,10,14,15^ Further studies utilizing precise quantitative methods would be needed to begin to understand the interaction between these bacterial species and other microbiota constituents.

The reduction in CAH observed clinically was not appreciated histologically as all Cb-infected mice, regardless of stock or the geographical site from which the mice were obtained, demonstrated histopathologic features of CAH and their total histopathology scores were higher than their respective uninfected controls. However, it is important to recognize that all the mice, except 1 which had a low clinical score at euthanasia, that had developed clinical disease were euthanized and their skin evaluated histologically after their clinical disease had resolved. The study design did not allow a comparison of histologic lesions between stocks at the point at which mice would be expected to have more severe dermatopathology and the differences among stocks, as well as controls, would likely have been markedly different. While the mean scores across Cb-infected mice ranged from 2.86 to 4.61 when euthanized at 14 dpi, as a comparison Cb-infected nude mice euthanized at the point in which their clinical scores were 2 and 3 had total histopathologic scores 3 to 4 times higher than the corresponding study mice that were sacrificed on 14 dpi (data not shown).

Ca colonization reflected in the total histopathology scores for A1 (Ca+) were significantly higher than the scores assigned to both stock B (Ca+) as well as C (Ca free) mice at 14 dpi suggesting that Ca colonization does not necessarily correlate with the dermatopathology severity observed when evaluated at this timepoint. Interestingly, there were minimal differences in total histopathology scores between Cb-infected mice evaluated at 14 or 28 dpi. Unfortunately, there is only a single study to the authors’ knowledge in which the long-term dermatopathology associated with Cb infection in nude mice was evaluated. The authors assessed histopathology of Cb-infected nude mice 12 weeks post infection.^7^ Significant histopathologic differences between Cb-infected and uninfected controls were not noted at this timepoint, but they did not compare the changes to earlier timepoints.^7^ It is unclear as to whether skin histology eventually returns to normal or resolves further as compared to our observations at 14 or 28 dpi. Clearly, the 14-day difference in the 2 stock B mouse groups evaluated is insufficient to result in histologic changes resulting from Cb infection. Future studies should characterize the long-term dermatopathology associated with Cb infection. Furthermore, while histopathology was appreciated in all Cb-infected mice, the degree of pathology was minimal to mild across stocks, likely implying a low impact to the overall health of the animals. However, this does not necessarily translate to its impact on research, especially on studies of the skin.

When assessing the scores of the individual histopathology criteria, the Cb-infected mice demonstrated significantly higher scores as compared to their respective uninfected controls. However, when comparing total histopathology scores between Cb-infected mice and their uninfected controls, significance was only appreciated in stock B mice. This discrepancy likely reflects the small number of control mice utilized.

In summary, this study demonstrated the potential impact of host genetics and/or the skin microbiota on the development and severity of CAH following Cb infection. Although there were notable differences observed, future studies are necessary to better understand the reasons for these differences with the goal of better understanding the biology of Cb, its impact on its’ immunocompromised host and ideally, eliminating or minimizing its’ impact in enzootically infected colonies.

## Abbreviations

Cb: *Corynebacterium bovis*.
Ca: *Corynebacterium amycolatum*.
CAH: *Corynebacterium*-associated hyperkeratosis.
dpi: days-post-infection.
CFU: Colony forming unit.
MPCS: Mean peak clinical score.

## Acknowledgements

We would like to acknowledge Kate Fodor, Glory Leung, Noah Mishkin, Michael Palillo, Juliette Wipf and the staff from the Laboratory of Comparative Pathology for their assistance with various technical components of the study.

